# ReCellTy: Domain-specific knowledge graph retrieval-augmented LLMs workflow for single-cell annotation

**DOI:** 10.1101/2025.04.23.650201

**Authors:** Dezheng Han, Yibin Jia, Ruxiao Chen, Wenjie Han, Shuaishuai Guo, Jianbo Wang

## Abstract

To enable precise and fully automated cell type annotation with large language models (LLMs), we developed a graph-structured feature–marker database to retrieve entities linked to differential genes for cell reconstruction. We further designed a multi-task workflow to optimize the annotation process. Compared to general-purpose LLMs, our method improves human evaluation scores by up to 0.21 and semantic similarity by 6.1% across 11 tissue types, while more closely aligning with the cognitive logic of manual annotation.

---

In single-cell RNA sequencing analysis, achieving precise cell type annotation through manual labeling typically requires two key steps: annotators retrieve relevant marker genes and integrate this information with their domain expertise to make informed decisions. Although various automated approaches have been developed, fully automated and precise cell type annotation remains a significant challenge. Prior to the advent of LLMs, traditional approaches (Fig. 1c), such as clustering analysis and feature matching, were applied for automated cell type annotation [1][2][3]. With the advancement of LLMs, their potential for fully or semi-automated cell annotation has also been explored (Fig. 1b) [4].

**Fig. 1.**
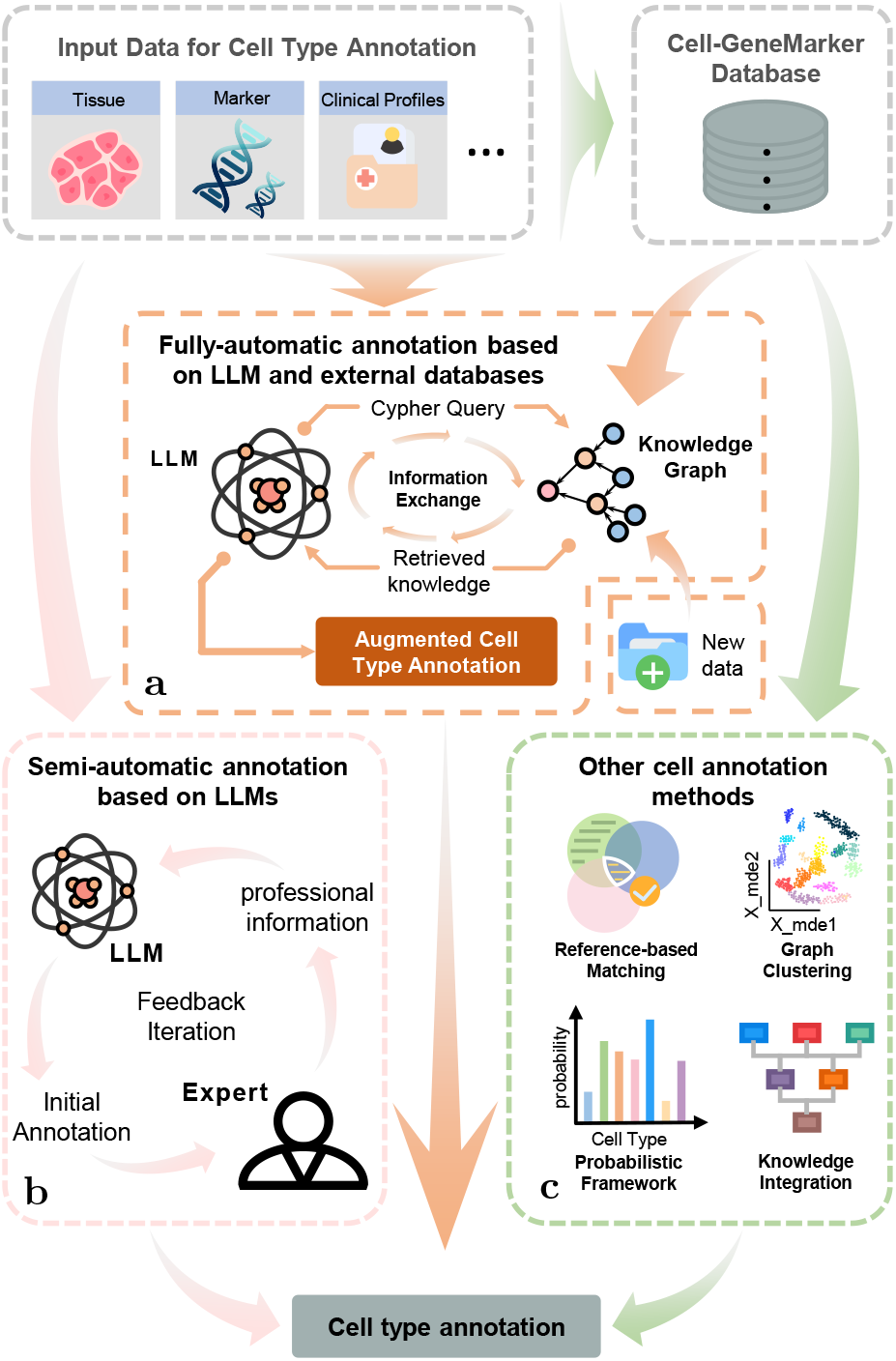
Overview of cell type annotation methods. **a**, Knowledge graph-driven LLM for automated annotation. **b**, Semi-automated annotation with expert-LLM collaboration. **c**, Traditional annotation methods prior to the advent of LLM.

However, general-purpose LLMs, not optimised for specific downstream tasks during pretraining, exhibit suboptimal performance in specialised domains. Existing approaches have attempted to fine-turn LLMs for cell annotation [5], but such methods often fail to fully capture the intricate relationships between genes and cell types, while continuous fine-tuning can result in catastrophic forgetting of previously learned data. To address these challenges, we propose equipping LLMs with a specialised knowledge graph and leveraging retrieval-augmented mechanisms to enhance performance in cell type annotation tasks (Fig. 1a).

Graph Retrieval-Augmented Generation (GraphRAG), an enhancement technique for LLMs introduced by Microsoft Research [6], enables LLMs to extract entities and their relationships from unstructured text. It constructs a structured knowledge graph and utilizes retrieval mechanisms to enhance its comprehension of entity relationships. In the medical domain, this method has been proposed [7], and its effectiveness has been demonstrated [8][9][10].In biological research, although existing studies have employed external data to enhance LLMs’ biological insights [11], knowledge graph construction has not been utilised to improve the retrieval and comprehension of biological entity information.

A potential limitation contributing to this phenomenon may stem from GraphRAG’s requirement for task-specific, high-quality datasets. We attempted to construct a knowledge graph using the raw CellMarker2.0 dataset [12], but the LLM still struggled to accurately identify the most probable cell type due to an overwhelming number of candidates. To address the data problem, we reconstructed the CellMarker2.0 database to make it more annotation-oriented. Additionally, we designed a multi-task workflow that simulates the manual annotation process, which systematically improves the annotation quality.

During the database reconstruction process, we decomposed cell naming conventions based on features, separating cell names into features and broad cell types while establishing explicit associations with relevant marker genes (Fig. 2c). Using this approach, we structurally reconstructed two datasets from the CellMarker2.0 database labeled as ‘Human’, encompassing over 78,000 raw entries. The final dataset consists of 1,528 naming features and 61,049 feature-gene-cell type associations (detailed data are available in Supplementary Table 1).

**Fig. 2.**
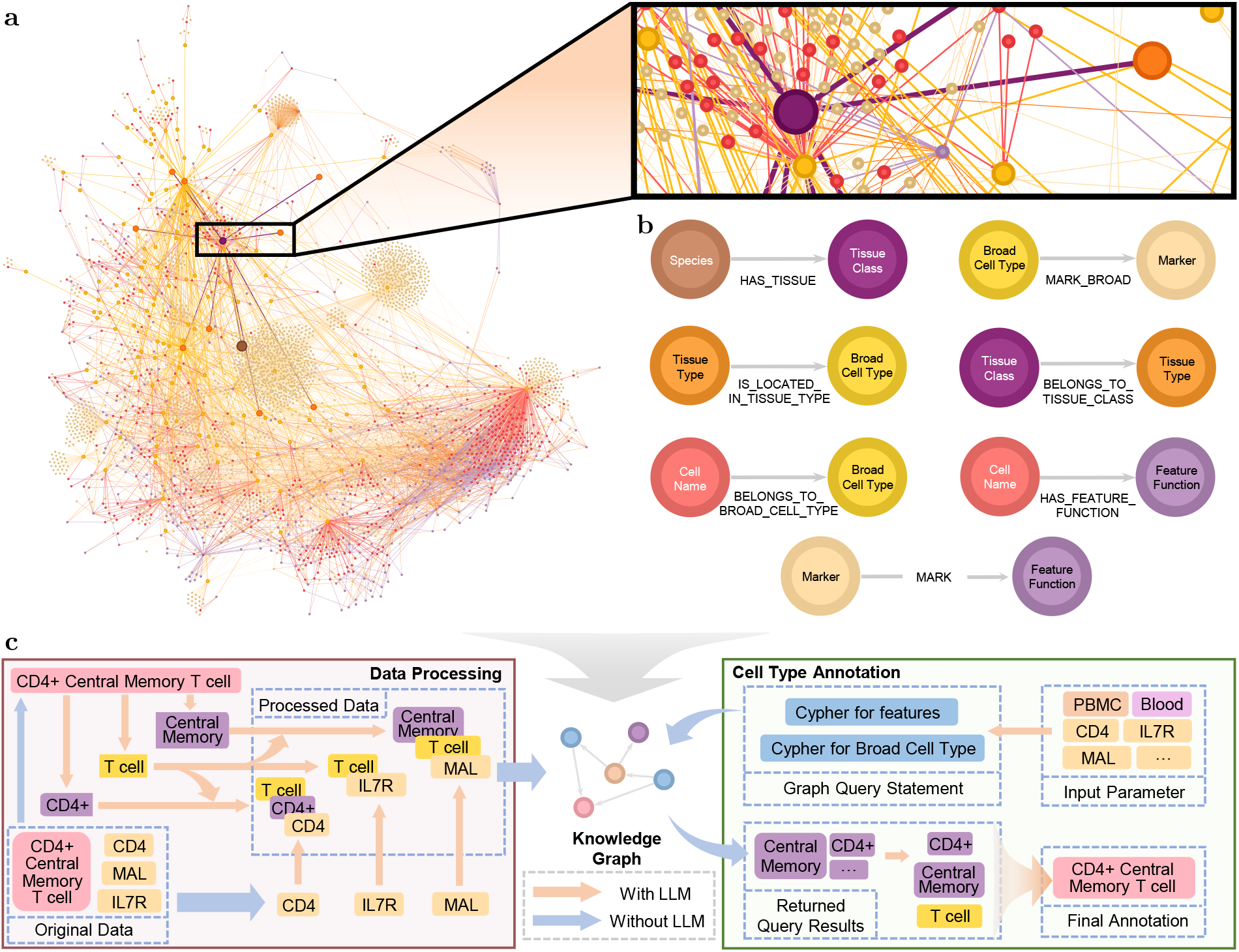
Structure and method of the knowledge graph-based cell type annotation framework. **a**, Visual representation of a subset of the data within the graph database. **b**, Seven node types and their relationship types within the graph database. **c**, Data processing pipeline and cell annotation question-answering workflow.

Notably, by leveraging the knowledge embedded within LLMs, we automated the data processing flow, eliminating the reliance on manual analysis. This approach transforms traditionally time-consuming and inefficient manual tasks into a scalable, high-throughput data processing pipeline. Furthermore, our method is adaptable to new datasets: By inputting gene and cell type information, the LLM-based framework rapidly parses, extracts biological data, and stores them in a structured format that is directly compatible with the knowledge graph, which significantly accelerates the integration of biological knowledge. Based on the restructured dataset, we constructed a structured knowledge graph by defining seven types of biological entity nodes and seven relationship types (Fig.2b), which were then stored in a graph database to facilitate efficient retrieval. This graph encompasses 18,850 biological entity nodes, including genes, cell types, and their associated features and functions, etc, as well as 48,944 connecting relationships between these nodes. The highly intricate structure tightly integrates various types of biological information, enabling the LLMs to uncover biological associations and interactions akin to human-level insights. We also visualised a subset of the database to provide an intuitive representation of the graph structure (Fig.2a).

The cell type annotation workflow is regarded as a cell name reconstruction process. We named our project ‘ReCellTy’, which stands for “Reconstructing Cell Types. In this process, the LLMs generate cypher code, a language used for querying graph database, to extract features and broad cell types related to differentially expressed marker genes from the database. These features and broad cell types are then organized and filtered by various agents to determine the most probable cellular signature, with the final agent performing the definitive cell type annotation based on the selected information (Fig. 2c). This modular restructuring enhances the flexibility of our pipeline in cell type annotation, while enabling detection of rare cell types that are often overlooked.

To demonstrate the effectiveness of our method, we conducted tests on 11 tissue types from the Azimuth dataset using four non-reasoning large language models. Given that the Claude 3.7 Sonnet model supports both standard and extended reasoning modes, we uniformly adopted the standard mode for all experiments in this study. For reproducibility, we performed five independent question-answering iterations for each row of differential genes, and the most frequent result was taken as our experimental evaluation benchmark (detailed annotation results are available in Supplementary Table 2).

Regarding the scoring mechanism design, we employed both manual evaluation and semantic evaluation scoring strategies to compare the annotation results of ReCellTy with the manual annotations in the dataset. Since our method supports the visual display of relevant information retrieved from the graph dataset, we introduced an intermediate process adjustment during manual evaluation to refine the accuracy of the final cell type scores. Semantic evaluation was assessed using an embedding model, where text annotations were converted into vectors, and the cosine similarity between them was calculated.

The experimental results demonstrate better performance of ReCellTy compared to general-purpose LLMs and the CellMarker 2.0 annotation across all four tested models and both scoring mechanisms (Fig. 3a)(Fig. 3d). After process adjustment, ReCellTy achieved an average score improvement of 0.18 across four models, with the most pronounced gain observed in deepseek-chat, reaching 0.21.

**Fig. 3.**
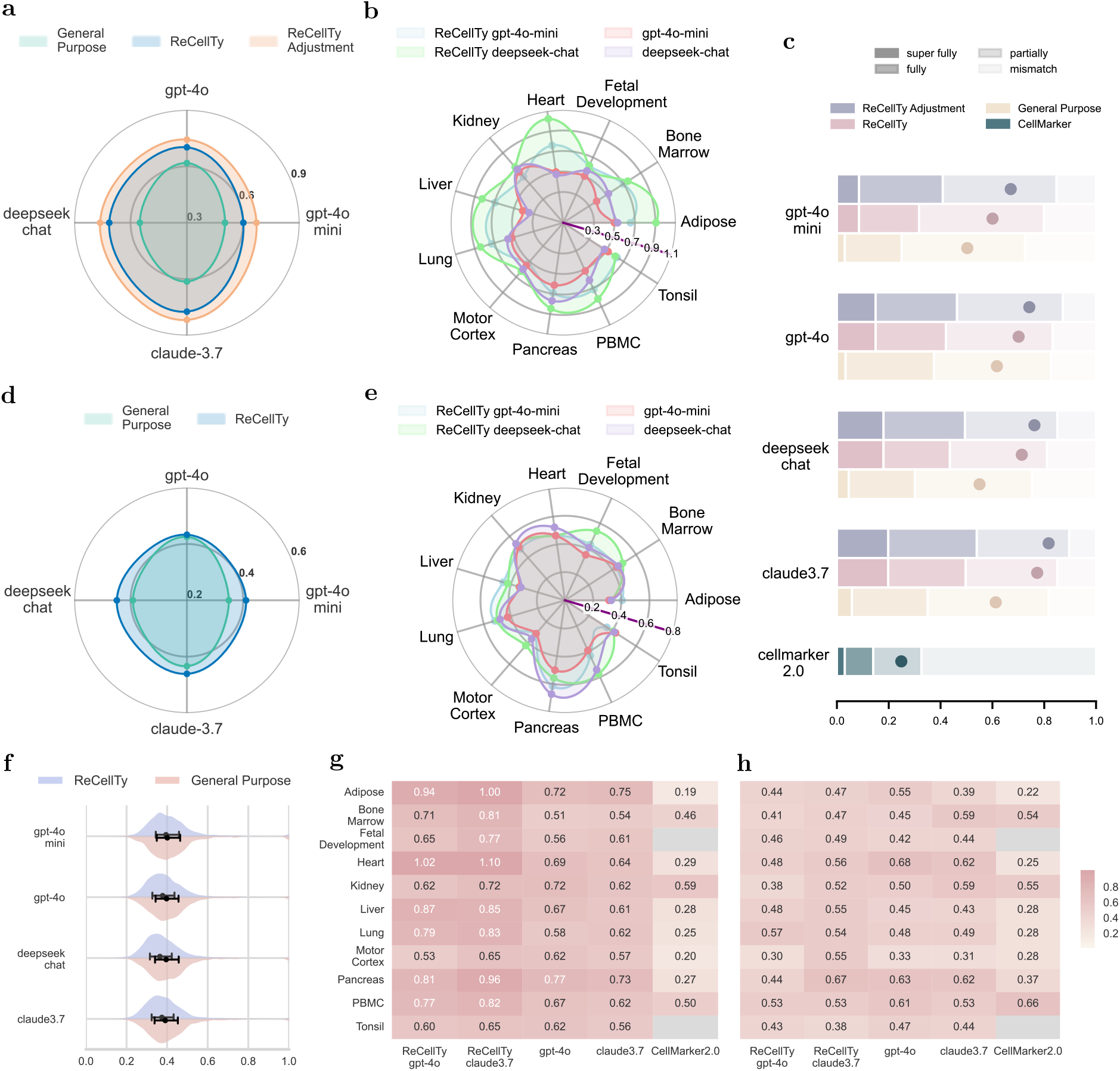
Performance evaluation. **a**, Human evaluation scores of four large models under different methods. **b**, Human evaluation scores of two large models across various tissues. **c**, Overall human evaluation scores for each method. **d**, Semantic evaluation scores of four large models under different methods. **e**, Semantic evaluation scores of two large models across various tissues. **f**, Intra-group semantic variance of annotation results for each model and method. **g**, Human evaluation scores of two models and CellMarker 2.0 across various tissues. **h**, Semantic evaluation scores of two models and CellMarker 2.0 across various tissues.

In manual evaluation, ReCellTy effectively bridges the performance gap between smaller and larger models (Fig. 3c). The general-purpose GPT-4o-mini model achieved a human evaluation score of only 0.50, but process adjustment ReCellTy boosted its performance to 0.67, surpassing larger models with more parameters and training data, including GPT-4o (0.62), DeepSeek-chat (0.55), and Claude 3.7 (0.61).

For each tissue, ReCellTy achieved higher manual evaluation scores in the majority of tissues (Fig. 3b)(Fig. 3g). For example, the deepseek-chat exhibited a 0.55 increase on the Heart dataset and a 0.48 improvement on the Liver dataset.

Although semantic evaluation generally reflects the semantic similarity between automated and manual annotations, we observed that it may assign lower scores to more specific cell type annotations. For example, in human evaluation, we assign a score of 1.5 when the model gives a more specific subtype, result shows that GPT-4o with ReCellTy reaches a score of 1.02, achieving a score improvement of 0.33 on the Heart dataset. This demonstrates that the model successfully annotated more specific cell sub-types. However, the semantic evaluation showed a 20% decrease. Although ReCellTy demonstrated overall improvement in semantic evaluation (average improvement: 3.8%; GPT-4o-mini: up to 6.1%), its ability to annotate more specific cell types, which should have been an advantage over other methods, instead resulted in less noticeable improvements in semantic scores at the level of individual tissues (Fig. 3e)(Fig. 3h).

To investigate the diversity of cell types generated by ReCellTy, we calculated the intra-group semantic similarity score for each model and method (Fig. 3f). The results showed that ReCellTy exhibited lower similarity compared to general-purpose large language models. This lower similarity indicates higher diversity, suggesting that by integrating knowledge graph information and employing retrieval-augmented generation. ReCellTy can focus on a more diverse range of cell types, significantly enhancing the diversity of annotation results compared to general-purpose LLMs.

Building upon these findings, future research may leverage LLM agents to autonomously manage and collaboratively utilise dynamic cell databases, enhancing their practical impact in cellular genomics.

## 1 Methods

### 1.1 Data processing

To systematically parse the associations between marker genes and cell features, we developed an end-to-end data processing pipeline based on the raw CellMarker2.0 database files (Cell marker Human.xlsx and Cell marker Seq.xlsx) [12]. Initially, raw data entries were categorised according to the ‘tissue class’ field, generating tissue-specific subsets. Within each subset, cells were further refined across four attribute dimensions, ‘cell name’, ‘cell type’, ‘cancer type’, and ‘tissue type’, to establish a hierarchical structure, where each distinct cell type was assigned a dedicated CSV (Comma-Separated Values) file. This multi-level decomposition strategy effectively isolates irrelevant data, significantly enhancing the LLM’s ability to focus on feature-marker relationships for a single target cell type.

For each generated CSV file, we designed a structured prompt template that takes cell name and the full marker gene list (‘marker’ field and ‘symbol’ field) as input. The LLM was tasked with three core functions: parsing function and feature descriptions from cell names, identifying broad cell type classifications, and establishing explicit feature-marker mappings. The model outputs CSV-formatted data containing these fields, which we extracted from the text responses and horizontally merged with additional metadata from the original files, forming an enhanced cell feature-marker data. For example, in the case of CD4+ cytotoxic T cells, the marker gene CD4 is directly linked to the naming feature CD4+, and since CD4 is biologically associated with T cells, we further mapped CD4 to T cell with CD4+ feature. This pipeline systematically covered all raw records for human species in CellMarker2.0, ensuring the completeness and traceability of feature association relationships.

Finally, the processed results were first aggregated vertically by tissue class into local datasets, before being globally integrated into a unified feature-marker association CSV database. In parallel, we developed an excel to enable intuitive data visualization.

### 1.2 Feature-marker GraphRAG

Based on feature-marker association data, we constructed a structured knowledge graph using the Neo4j graph database https://neo4j.com/. Specifically, we developed a Cypher query code to sequentially convert preprocessed CSV data into graph entities. We defined seven core node types, including Marker (gene marker) and FeatureFunction (features or functions in cell name), and established seven types of semantic relationships, such as MARK (annotation from Marker to FeatureFunction) and HAS FEATURE FUNCTION (annotation from CellName to FeatureFunction), forming a multi-layered cellular feature network.

For retrieval, we leveraged the Graph-CypherQAChain module from the LangChain https://www.langchain.com, integrating retrieval and question-answering into a high-efficiency querying system. The core mechanism involves feeding the LLM a schema description of the Neo4j graph, reinforcing its understanding of the knowledge graph’s topology and improving the quality of Cypher query generation. The query workflow proceeds as follows: first, the LLM parses the user’s natural language input to determine key retrieval elements. Second, a Cypher query is dynamically generated based on the knowledge graph’s structural features. Third, the query is executed to retrieve structured data from Neo4j. Finally, the retrieval results are combined with the original query for final inference, generating a formatted response.

For cell type annotation, we implemented two key optimisations in the GraphCypherQAChain framework. First, we introduced cell type annotation specific prompt templates, embedding example Cypher queries as references to improve generation accuracy. Second, we designed a dual-retrieval augmentation mechanism, where the system first retrieves broad cell type for given markers, ensuring a category-level prediction, and in parallel, the system retrieves functional feature descriptions linked to the same markers, constructing a structured feature mapping table. The standardised outputs from these two retrieval pathways serve as input evidence for the down-stream agent-based decision-making system.

### 1.3 Multi-Task framework

To enable human-like decision logic in cell type annotation, we developed a modular workflow system built on top of the knowledge graph retrieval architecture. This workflow integrates the dual-retrieval augmentation mechanism (broad cell type retrieval and feature-function mapping) and employs a multi-task system to refine knowledge-driven inference. Specifically, the workflow consists of the following components:

**CellType Query Task** processes input top differentially expressed genes to retrieve associated broad cell types from the graph dataset, summarizing results in a standardized Marker-CellType correspondence format.

**CellType Selection Task** makes determinations about broad cell type based on the summarized information from the CellType Query Task.

**Feature Query Task** functions similarly to the CellType Query Task but returns associated named features.

**Feature Selection Task** filters the retrieved marker–feature mappings, selecting 2–3 high-probability features to reduce decision space complexity for downstream processing.

**CellType Annotation Task** integrates multiple dimensions of information, including the predicted broad cell type, the selected features, and the original marker genes to label the final cell type annotation.

The prompts for each task, including the data processing task, are provided in the Supplementary Prompt.

### 1.4 Comparative methods

The method proposed in this study achieves enhanced cell type annotation performance by integrating and optimizing LLMs with the knowledge augmentation strategy of the CellMarker 2.0 database. To validate the effectiveness of our method, we selected the following two methods for comparative experiments:

**CelltypeGPT**. This is an LLM-based cell annotation tool [4]. Since ReCellTy employs a single-row differentially expressed gene processing mode, to ensure fairness in the comparison, we adjusted the prompting strategy of CelltypeGPT and denote its method as ‘general-purpose’. The adjusted prompt template is as follows:

’Identify cell types of TissueName cells using the following markers. Only provide the cell type name. Do not show numbers before the name. Some can be a mixture of multiple cell types. GeneList’

TissueName and GeneList will be replaced with the actual tissue and differentially expressed gene list, respectively.

**CellMarker2.0**. The official tool, directly accessing the webpage interface of this database for annotation.

### 1.5 Evaluations

#### Manual evaluation

For manual evaluation, we improved upon the evaluation framework of CelltypeGPT. The criteria for determining the final annotation results are as follows: if the automatically annotated result is more specific than the manual annotation, it is defined as “super fully”; if the two are exactly the same, it is “fully”; if the two belong to the same major cell type or have a differentiation-related connection, it is defined as “partially”; and completely unrelated is “mismatch”. In the intermediate process evaluation, if the major cell type and the selected relevant features can be combined to produce a cell type that is exactly the same as the manual annotation, it is “fully”; partially related is “partially”; and completely unrelated is “mismatch”. The four evaluation levels (super fully/fully/partially/mismatch) are assigned weights of 1.5, 1.0, 0.5, and 0, respectively, and then the average score within the group is calculated.

#### Semantic evaluation

For semantic similarity, we used OpenAI’s text-embedding-3-small to encode the cell annotation results and then calculated the cosine similarity between the vector representations of the manual and automatic annotations. Subsequently, we normalized the cosine similarity within the group and divided it into five equal intervals from high to low, assigning scores of 1, 0.75, 0.5, 0.25, and 0, respectively, and then calculated the average score within the group.

### 1.6 Application

To enhance system interpretability and assist user decision-making, we developed a multi-stage visualisation system and an interactive user interface (UI). The core UI modules include:

**Input Layer**, allowing users to submit top differential marker genes and select tissue. The selection of specific tissues aims to narrow the query and evaluation scope, thereby improving accuracy. To address data scarcity in certain tissues, we also introduced a global query mode that removes retrieval constraints, allowing comprehensive utilization of the graph data.

**Processing Layer**, displaying real-time workflow status (retrieval progress and agent decisionmaking steps).

**Output Layer**, presenting final annotation results alongside intermediate reasoning steps, including broad cell type and features selection. This UI provides an end-to-end transparent pipeline, allowing users to trace the annotation process from raw input data to final cell type predictions.

Additionally, to ensure seamless integration with single-cell sequencing analysis pipelines (e.g., Seurat), we developed a Python-based package for processing the top differential expressed genes. Although our package operates in a Python environment, compatibility with upstream workflows is enabled through cross-language interoperability packages rpy2 (allows calling R from Python) and reticulate (allows calling python from R).

## Supporting information

Supplementary Prompt

## 2 Data availability

The datasets and models used in this study are available from the following sources. The Cell-Marker 2.0 dataset can be downloaded at http://www.bio-bigdata.center/CellMarker_download.html, and the Azimuth dataset is available at https://azimuth.hubmapconsortium.org/. The language models employed, including GPT-4o-mini, GPT-4o, and text-embedding-3-small, are accessible via the OpenAI API at https://openai.com/api/. DeepSeek-chat and Claude 3.7 were accessed through their respective APIs at https://platform.deepseek.com/sign in and https://www.anthropic.com/api.

## 3 Code availability

The ReCellTy package and UI, together with their source code and associated data, are publicly available at https://github.com/SSG2019/ReCellTy. A portion of the original code framework, data processing pipeline, and experimental test datasets has also been released at https://github.com/SSG2019/ReCellTy-paper, with the remaining components to be made available upon acceptance of the manuscript.

## References

[1] Ianevski, A., et al.: Fully-automated and ultra-fast cell-type identification using specific marker combinations from single-cell transcriptomic data. Nat. Commun. 13, 1246 (2022) 10.1038/s41467-022-28803-w

[2] Aran, D., et al.: Reference-based analysis of lung single-cell sequencing reveals a transitional profibrotic macrophage. Nat. Immunol. 20, 163–172 (2019) 10.1038/s41590-018-0276-y

[3] Xu, J., et al.: CIForm: a transformer-based model for cell-type annotation of large-scale single-cell RNA-seq data. Brief. Bioinform. 24(4), 195 (2023) 10.1093/bib/bbad195

[4] Hou, W., et al.: Assessing GPT-4 for cell type annotation in single-cell RNA-seq analysis. Nat. Methods 21, 1462–1465 (2024) 10.1038/s41592-024-02235-4

[5] Levine, D., et al.: Cell2Sentence: teaching large language models the language of biology. bioRxiv (2023) 10.1101/2023.09.11.557287

[6] Darren, E., et al.: From local to global: A graph RAG approach to query-focused summarization (2025) 2404.16130 [cs.CL]

[7] Gilbert, S., et al.: Augmented non-hallucinating large language models as medical information curators. npj Digit. Med. 7, 100 (2024) 10.1038/s41746-024-01081-0

[8] Wu, J., et al.: Medical Graph RAG: Towards safe medical large language model via graph retrieval-augmented generation (2024) 2408.04187 [cs.CV]

[9] Zuo, K., et al.: KG4Diagnosis: A hierarchical multi-agent LLM framework with knowledge graph enhancement for medical diagnosis (2025) 2412.16833 [cs.AI]

[10] Zhao, X., et al.: MedRAG: Enhancing retrieval-augmented generation with knowledge graph-elicited reasoning for healthcare copilot (2025) 2502.04413 [cs.CL]

[11] Liu, W., et al.: DrBioRight 2.0: an LLM-powered bioinformatics chatbot for large-scale cancer functional proteomics analysis. Nat. Commun. 16, 2256 (2025) 10.1038/s41467-025-57430-4

[12] Hu, C., et al.: Cellmarker 2.0: an updated database of manually curated cell markers in human/mouse and web tools based on scRNA-seq data. Nucleic Acids Res. 51(D1), 870–876 (2022) 10.1093/nar/gkac947

