## Supplementary Prompt for "ReCellTy: Domain-specific knowledge graph retrieval-augmented LLMs workflow for single-cell annotation"

This document is the supplementary information for the agent prompt of the paper "ReCellTy: Domain-specific knowledge graph retrieval-augmented LLMs workflow for single-cell annotation".

### 1 Data Processing Task

##### system prompt

```
You are a professional bioinformatics analyst specializing in handling CSV files, especially in
the field of cells and genes.
your task is to analyze the task and output the required cvs file based on the given content.
```

##### user prompt

```
Your response should be as follows:
Firstly, separate the features and functions in the cell name.
Secondly, Analyze the relationship between each gene and feature&function, and save the genes
that can reflect the feature function into cvs file species.
If the input cell itself belongs to a major cell type without any feature or function, all genes
will be output for each row, and the feature & function fields will be left blank.
The output cvs file header includes: broad_cell_types, feature&function, marker
note: The markers in each line of the generated cvs file should determine the feature&function.
cell name: {cell_name},
marker: {marker}(Please analyze every marker, some genes may be associated with traits, while
others are not).

Here is an example:
input:
cell name: Effector CD4+ T cell
marker: markerA, markerB, markerC, markerD, markerE,
output:
markerA is associated with CD4+ ,markerE is associated with Effector.
'''csv
broad_cell_types,feature&function,marker
T cell,CD4+,markerC
T cell,CD4+,markerA
T cell,Effector,markerE
T cell,Effector,markerB
T cell,Effector,markerD
'''
```

'cell_name' and 'marker' are replaced with specific cell names and marker genes during data processing procedure.

### 2 CellType Query Task

#### 2.1 Task for cypher generation

##### user prompt (specific tissues)

```
Task:Generate Cypher statement to query a graph database.
Instructions:
Use only the provided relationship types and properties in the schema.
Do not use any other relationship types or properties that are not provided.
Schema:
{schema}
Note: Do not include any explanations or apologies in your responses.
Do not respond to any questions that might ask anything else than for you to construct a
Cypher statement.
Do not include any text except the generated Cypher statement.
Examples: Here are a few examples of generated Cypher statements for particular questions:
Note: only change the marker and 'tissue_class' according to the question, do not change the
rest of the Cypher code.
MATCH (m:Marker)-[:MARK_BROAD]->(bct:BroadCellType)-[:IS_LOCATED_IN_TISSUE_TYPE]->(t: TissueType)
-[:BELONGS_TO_TISSUE_CLASS]->(tc: TissueClass)
WHERE m.name IN ['Marker1', 'Marker2', 'Marker3'...]
  AND tc.name = 'tissue_class'
RETURN m.name AS Marker, COLLECT(DISTINCT bct.name) AS BroadCellType


The question is:
{question}
```

##### user prompt (global)

```
Task:Generate Cypher statement to query a graph database.
Instructions:
Use only the provided relationship types and properties in the schema.
Do not use any other relationship types or properties that are not provided.
Schema:
{schema}
Note: Do not include any explanations or apologies in your responses.
Do not respond to any questions that might ask anything else than for you to construct a
Cypher statement.
Do not include any text except the generated Cypher statement.
Examples: Here are a few examples of generated Cypher statements for particular questions:
Note: only change the marker according to the question, do not change the rest of the
Cypher code.
MATCH (m:Marker)-[:MARK_BROAD]->(bct:BroadCellType)
WHERE m.name IN ['Marker1', 'Marker2', 'Marker3'...]
RETURN m.name AS Marker, COLLECT(DISTINCT bct.name) AS BroadCellType


The question is:
{question}
```

The prompt is provided by LangChain, and we have modified the query examples. In actual applications, 'schema' will be replaced by the description architecture of the graph database, and 'question' will be replaced by the user prompt of '2.2 Task for summary information'.

#### 2.2 Task for summary information

##### user prompt (specific tissues)

```
I need to perform cell type annotation based on given markers.
Your task is to summarize the broad cell types corresponding to the relevant genes.
Markers: {marker}
TissueClass: {tissue}
Summary template is as follows:
marker1: BroadCellType1, BroadCellType2, BroadCellType3....
marker2: BroadCellType1, BroadCellType2, BroadCellType3....
.....
```

##### user prompt (global)

```
I need to perform cell type annotation based on given markers.
Your task is to summarize the broad cell types corresponding to the relevant genes.
Markers: {marker}
Summary template is as follows:
marker1: BroadCellType1, BroadCellType2, BroadCellType3....
marker2: BroadCellType1, BroadCellType2, BroadCellType3....
.....
```

The 'marker' will be replaced by the input top differentially expression genes, and the 'Tissue' will be replaced by the input tissue class.

### 3 CellType Selection Task

##### system prompt

```
You are an expert in the field of biological cell marker functions. Your task is to perform a
step in cell type annotation: identify broad cell types based on relevant information and
tissue types.
```

##### user prompt

```
The information provides possible major cell types: {broad_cell_type_info}
The tissue is {Tissue}
Give me your final judgment(The only one broad type of cell).
Return only the cell type as the output.
```

The 'broad_cell_type_info' will be replaced by the output of '2.2 Task for summary information', and 'Tissue' will be replaced by the input tissue type.

### 4 Feature Query Task

#### 4.1 Task for cypher generation

##### user prompt (specific tissues)

```
Task:Generate Cypher statement to query a graph database.
Instructions:
Use only the provided relationship types and properties in the schema.
Do not use any other relationship types or properties that are not provided.
Schema:
{schema}
Note: Do not include any explanations or apologies in your responses.
Do not respond to any questions that might ask anything else than for you to construct a
Cypher statement.
Do not include any text except the generated Cypher statement.
Examples: Here are a few examples of generated Cypher statements for particular questions:
Note: only change the marker and 'tissue_class' according to the question, do not change the
rest of the Cypher code.
MATCH (m:Marker)-[:MARK]->(ff:FeatureFunction)<-[:HAS_FEATURE_FUNCTION]-(cn:CellName)-
[:BELONGS_TO_BROAD_CELL_TYPE]->(bct:BroadCellType)
-[:IS_LOCATED_IN_TISSUE_TYPE]->(t: TissueType)-[:BELONGS_TO_TISSUE_CLASS]->(tc: TissueClass)
WHERE m.name IN ['Marker1', 'Marker2', 'Marker3'...]
  AND tc.name = 'Tissue_class'
RETURN m.name AS Marker, COLLECT(DISTINCT ff.name) AS FeatureFunctions


The question is:
{question}
```

##### user prompt (global)

```
Task:Generate Cypher statement to query a graph database.
Instructions:
Use only the provided relationship types and properties in the schema.
Do not use any other relationship types or properties that are not provided.
Schema:
{schema}
Note: Do not include any explanations or apologies in your responses.
Do not respond to any questions that might ask anything else than for you to construct a
Cypher statement.
Do not include any text except the generated Cypher statement.
Examples: Here are a few examples of generated Cypher statements for particular questions:
Note: only change the marker according to the question, do not change the rest of the
Cypher code.
MATCH (m:Marker)-[:MARK]->(ff:FeatureFunction)
WHERE m.name IN ['Marker1', 'Marker2', 'Marker3'...]
RETURN m.name AS Marker, COLLECT(DISTINCT ff.name) AS FeatureFunctions


The question is:
{question}
```

The prompt is provided by LangChain, and we have modified the query examples. In actual applications, 'schema' will be replaced by the description architecture of the graph database, and 'question' will be replaced by the user prompt of '4.2 Task for summary information'.

#### 4.2 Task for summary information

##### user prompt (specific tissues)

```
Query and summarize the feature functions of the following markers.
Markers: {marker}
TissueClass: {tissue}
Summary template is as follows:
marker1: feature1, feature2, feature3....
marker2: feature1, feature2, feature3....
.....
```

##### user prompt (global)

```
Query and summarize the feature functions of the following markers.
Markers: {marker}
Summary template is as follows:
marker1: feature1, feature2, feature3....
marker2: feature1, feature2, feature3....
.....
```

The 'marker' will be replaced by the input top differentially expression genes, and the 'Tissue' will be replaced by the input tissue class.

### 5 Feature Selection Task

##### system prompt

```
You are an expert in the field of biological cell marker functions. Your task is to perform
a step in cell type annotation.
Based on the determined broad cell type and the feature functions corresponding to the markers,
select at most three features/functions that you believe are most likely to be expressed.
Please first analyze according to the requirements.
```

##### user prompt

````
The broad cell type is known: {broad_type}.
the corresponding feature functions for each marker are as follows: {marker_feature}
Select the relevant features according to the following three steps:
First, analyze the correlation between other feature functions and markers, and make
the selection.
Second, check if there are feature function names that highly overlap with the marker name.
If so, the feature/function must be included in your answer.
for example, if a marker gene is named 'ABC' and its corresponding features include 'ABC+',
then 'ABC+' is a mandatory option.
Finally, check the number of features corresponding to each marker. If a marker corresponds to
only one or two feature functions. If so, the feature/function must be included in your answer.
In ang cases, your answer should include a answer string, e.g.:
```answer
feature_one, feature_two, feature_three
```
Do not include markers in the answer string
````

The 'broad_type' will be replaced by the output of '3 CellType Selection Task'. The 'marker_feature' will be replaced by the output of '4.2 Task for summary information',

### 6 CellType Annotation Task

##### system prompt

```
You are an intelligent assistant tasked with annotating cell types.
Given the determined broad cell type and the associated features and functions,
determine the most likely cell type.
```

##### user prompt

```
Given the following information:
marker: {marker}
broad cell type: {broad_cell}
feature&function: {feature_function}

Return only the cell type as the output.
```

The 'marker' will be replaced by the input top differentially expression genes, 'broad_cell' will be replaced by the output of '3 CellType Selection Task', and 'feature_function' will be replaced by the output of '5 Feature Selection Task'.
